# Consistent ultra-long DNA sequencing with automated slow pipetting

**DOI:** 10.1101/2020.09.18.302471

**Authors:** Trent M. Prall, Emma K. Neumann, Julie A. Karl, Cecilia G. Shortreed, David A. Baker, Hailey E. Bussan, Roger W. Wiseman, David H. O’Connor

## Abstract

**Background:** Oxford Nanopore Technologies instruments can sequence reads of great length. Long reads improve sequence assemblies by unambiguously spanning repetitive elements of the genome. Sequencing reads of significant length requires the preservation of long DNA template molecules through library preparation by pipetting reagents as slowly as possible to minimize shearing. This process is time-consuming and inconsistent at preserving read length as even small changes in volumetric flow rate can result in template shearing.

**Results:** We have designed SNAILS (Slow Nucleic Acid Instrument for Long Sequences), a 3D-printable instrument that automates slow pipetting of reagents used in long read library preparation for Oxford Nanopore sequencing. Across six sequencing libraries, SNAILS preserved more reads exceeding one hundred kilobases in length and increased its libraries’ average read length over manual slow pipetting.

**Conclusions:** SNAILS is a low-cost, easily deployable solution for improving sequencing projects that require reads of significant length. By automating the slow pipetting of library preparation reagents, SNAILS increases the consistency and throughput of long read Nanopore sequencing.

## Background

Oxford Nanopore Technologies’ (ONT) MinION and GridION instruments generate reads of unprecedented length [1]. Long reads improve genomic assemblies by spanning repetitive regions and structural variation within the genome. These regions have historically complicated alignments created solely using shorter, second-generation sequencing reads [2]. An increased understanding of these genomic features is crucial as evidence linking heritable structural variation with disease phenotypes is emerging [3]. ONT sequencing has resolved the nucleotide identity of gaps left by 100 base pair (bp) Illumina reads aligned to GRCh38 better than other long read sequencing technologies [4]. These gaps included coding exons for 76 genes with known disease-associated mutations cataloged in the Human Gene Mutation Database [5]. A pathogenic deletion in TMEM231, a gene known to be involved in Meckel-Gruber syndrome, was uncovered by ONT reads [6]. This mutation had been masked by ambiguous mapping of short reads to a paralogous pseudogene locus. Recently, the first telomere-to-telomere assemblies of the human X and Y chromosomes were created using ONT sequencing [7, 8]. In these assemblies, ultra-long reads in excess of one megabase pair (Mb) were used to bridge the previous assemblies’ centromeric gaps. Using similar methods, ONT generated long reads have been utilized to establish more contiguous assemblies for important model organisms [9–12].

Sequencing long reads requires isolating high molecular weight (HMW) DNA and preserving those DNA molecules through library preparation [13]. Methods for obtaining and preserving HMW DNA have been developed for use in the field of optical mapping, which benefits from continuous strands of DNA [14]. For example, DNA in solution can be condensed into a compact conformation by adding neutral polymers that exert osmotic pressure to nearby double helices [15]. Condensed DNA can also be induced by adding cationic ligands into solution to promote attractive interactions of DNA segments. DNA is resistant to shearing by pipetting when in condensed conformation [16]. Unfortunately, elongation of condensed DNA requires monovalent salts that could interfere with ONT sequencing pores. In 1984, Schwartz et al. pioneered pulsed gradient gel electrophoresis that allowed for the separation of chromosomal-sized DNA [17]. Consistent ultra-long DNA isolation was accomplished by suspending cells in low-melting-point agarose and diffusing lysing reagents into the resulting suspension. The “nuclei-method” later expanded on this technique by enriching cell nuclei before embedding in agarose to remove cytoplasmic and other cellular lysates from the resulting DNA [18]. Second-generation short read sequencing can be performed on DNA extracted using the nuclei method [19]. However, gel extraction protocols designed for preserving high molecular weight DNA sequences require access to specialized electroelution chambers or pulsed-field gel electrophoresis systems [20]. HMW DNA can also be obtained using phenol-chloroform extractions based on a previously published protocol [21]. This protocol uses caustic chemicals and requires several hours to perform [13]. Alternatively, Circulomics Inc. has developed a thermoplastic substrate that can bind and release large quantities of DNA without damaging it [22]. They have created an isolation kit (Nanobind) using the substrate that produces large amounts of HMW DNA in approximately sixty minutes. Recently, Circulomics has reported a 2.44 Mb continuous read sequenced by MinION representing the longest single read ever sequenced [23]. Regardless of the isolation method, the resulting HMW DNA must undergo library preparation, which introduces more opportunities for shearing to occur.

Long read ONT libraries are prepared using the Rapid Sequencing Kit (SQK-RAD004). This protocol involves simultaneous fragmentation and ligation of tags by a transposome complex followed by the attachment of sequencing adaptors to template molecules before loading onto flow cells for sequencing. Throughout library preparation, DNA is susceptible to shearing by pipetting. A single draw using a normal-bore pipette tip shears megabase DNA, so wide-bore tips are used to reduce the amount of shearing [16]. DNA shearing can also be mitigated by pipetting at a slow rate [16]. Because the standard RAD004 sequencing protocol wasn’t initially intended for long read sequencing, a modified RAD004 protocol was developed to preserve template length [24]. In this protocol, HMW DNA and sequencing reagents are mixed by pipetting up and down as slowly as possible eighteen total times [25]. N50 denotes the minimum length for which half of the read lengths in a pool are equal to or longer, and it is used to estimate the distribution of read lengths within a pool. When implemented, the modified RAD004 protocol increased the average N50 from 10.6 kilobases across thirty-nine standard library preparations to 99.7 kilobases across two modified library preparations from a single sample [24]. Performing the long read RAD004 protocol for a single flow cell typically requires two hours of manual pipetting to perform [13]. This process can be a significant challenge for technicians to effictively execute and is limited in throughput. To address this challenge, we have designed and tested SNAILS (Slow Nucleic Acid Instrument for Long Sequences), an open-sourced, 3D-printed robot that automates slow pipetting steps used in long read RAD004 library preparations. SNAILS eliminates some of the variables inherent to manual slow pipetting caused by unsteady hands. Additionally, SNAILS dramatically increases the long read sequencing throughput capabilities of a single technician.

## Results

### SNAILS - Slow Nucleic Acid Instrument for Long Sequences

We hypothesized that slow pipetting reagents could be automated by rotating a pipette plunger using a DC motor. To accomplish this, we created SNAILS, a 3D-printed scaffold designed around the VistaLab Ovation M micropipette that aligns a twelve-volt direct current planetary gear motor with a three-axle gear train connected to the pipette’s plunger (Fig 1A, B). An L298N motor driver supplies current to the motor and is controlled through 3.3-volt signals generated by a raspberry pi 3B+ microcontroller. This configuration allows the direction of current, and by extension the direction of motion, to be programmatically toggled to aspirate or dispense volumes (Fig 1C). The three-axle gear train creates a 36:7 ratio between the motor and pipette plunger that downshifts the motor’s twelve rotations per minute to approximately 1.25 rotations per minute (Fig 1D). This results in a volumetric flow rate of around 6 μl per minute when used with a P100 micropipette. SNAILS are printed in multiple pieces that are assembled by connecting dovetailed joints attached to each piece (Supplementary Figures 1-4) Files for printing, assembly instructions, and source code for SNAILS are freely available for download from github (https://github.com/dholab/SNAILS).

**Figure 1.**
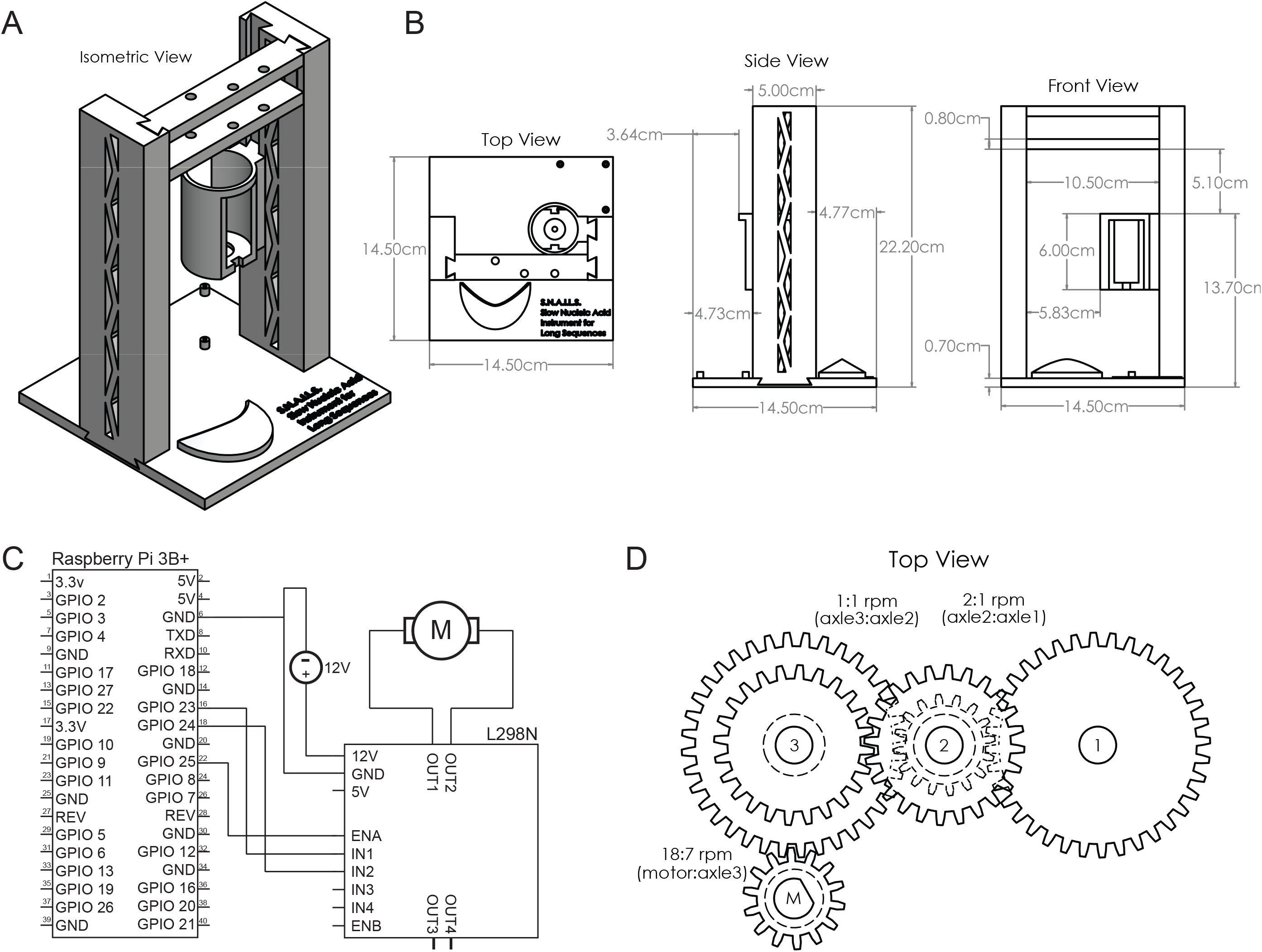
Slow Nucleic Acid Instrument for Long Sequences schematics. **A.** Rendering of a fully assembled SNAILS from five individual pieces. Each piece is printed with dovetail joints that are slid together during assembly. **B.** Dimensions of the fully assembled instrument. **C.** Circuit diagram for SNAILS motor, L298N, and microcontroller general-purpose input/output pins are representative of a Raspberry Pi 3B+ however, any microcontroller with configurable pins can be used. **D.** Rotations per minute ratios for four-axle gear train. M represents the motor’s axle.

### Comparison of high molecular weight DNA isolations

Three HMW DNA isolations were performed using approximately 2 × 10^7^ peripheral blood mononuclear cells per isolation from a single rhesus macaque. Eighteen sequencing libraries were prepared from these DNA isolations using ONT SQK-RAD004 library preparation kits. One of three pipetting methods was used for library preparation: manual aspiration using an Eppendorf P200 micropipette, manual rotation using an Ovation M P100 micropipette, and automated rotation of an Ovation M P100 micropipette using SNAILS. Manual aspirational and rotational pipetting were performed by technicians with considerable experience in long read RAD004 preparations. Rotational pipetting was performed by rotating the pipette’s plunger as slowly as possible to draw and dispense liquids. SNAILS were operated by more novice technicians with no experience performing long read library preparations by hand. Each of the pipetting methods was used to create two libraries simultaneously from each isolation to produce of six sequencing runs per pipetting method. It has been observed that MinKNOW arbitrarily fragments base called reads during sequencing [26]. Therefore, we informatically fused reads post-sequencing to accurately assess the read length retention between pipetting methods using whale_watch.py from the BulkVis toolset [26]. This process also ensures that all analyzed reads successfully map to the hg38 reference genome to eliminate the possibility of spurious long reads that have been observed during Nanopore sequencing runs (see Methods). The results of each library preparation are displayed in Table 1. We calculated the mean N50 across all three pipetting methods to estimate the relative length distribution. Future studies might benefit from quantification of HMW DNA using pulsed field gel electrophoresis, as HMW DNA is highly viscous and difficult to measure with simpler quantification methods. N50 values vary substantially per isolation with means of 23,124 bp; 15,741 bp; and 19,975 bp from isolations one, two, and three, respectively. All eighteen N50s fell below the expected range of greater than 50 kb. It should be noted that DNA extractions were performed from cryopreserved cells. DNA extracted from cryopreserved cells has been shown to yield slightly shorter lengths than DNA extracted from fresh cells [24]. This suggests that the input gDNA was not exceptionally HMW. However, a considerable amount of ultra-long reads greater than 100 kb were generated in the study, and so the pipetting methods still demonstrate long read retention. The length of DNA within a pool can vary substantially between HMW isolations. Therefore, DNA length is still limited by HMW isolations and automated library preparation serves only to preserve existing long molecules.

**Table 1.** Results of eighteen SQK-RAD004 long read sequencing libraries. Libraries were prepared in duplicate using one of three pipetting methods. All sequencing libraries were sequenced using Oxford Nanopore Technologies’ GridION instrument. The pores available at start of run were calculated by the MinKNOW software and recorded at the start of sequencing.

| DNA isolation | Method | Pores available at run start | Number of bases (Gb) | Number of reads | Longest read (bp) | N50 size (bp) | N90 size (bp) |
| --- | --- | --- | --- | --- | --- | --- | --- |
| 1 | Aspiration | 1505 | 4.25 | 661189 | 737462 | 21327 | 2212 |
|  | Aspiration | 1528 | 3.92 | 644261 | 645672 | 19617 | 2077 |
|  | Rotation | 1432 | 2.56 | 283481 | 862322 | 25482 | 3818 |
|  | Rotation | 1362 | 4.32 | 365489 | 782247 | 30174 | 4891 |
|  | Automatio | 1452 | 4.97 | 795794 | 1021308 | 13082 | 2333 |
|  | Automatio | 1442 | 4.92 | 496674 | 1253204 | 29063 | 3933 |
|  | Mean | 1454 | 4.16 | 541148 | 883703 | 23124 | 3211 |
| 2 | Aspiration | 883 | 3.34 | 673876 | 682744 | 12695 | 1909 |
|  | Aspiration | 1093 | 3.16 | 598310 | 774556 | 13510 | 2030 |
|  | Rotation | 1046 | 5.4 | 617128 | 508355 | 17107 | 3407 |
|  | Rotation | 1208 | 7.41 | 637349 | 526752 | 24163 | 4800 |
|  | Automatio | 1182 | 7.21 | 1209053 | 508355 | 12382 | 2342 |
|  | Automatio | 1571 | 10.92 | 1608258 | 526752 | 14589 | 2654 |
|  | Mean | 1164 | 6.24 | 890662 | 587919 | 15741 | 2857 |
| 3 | Aspiration | 1420 | 5.06 | 883721 | 605755 | 11809 | 2127 |
|  | Aspiration | 1629 | 6.46 | 755277 | 798240 | 18470 | 3381 |
|  | Rotation | 1311 | 8.02 | 915877 | 661883 | 16458 | 3566 |
|  | Rotation | 818 | 4.24 | 386642 | 584414 | 19942 | 4285 |
|  | Automatio | 1421 | 4.97 | 596368 | 602104 | 21358 | 3539 |
|  | Automatio | 1410 | 6.89 | 545627 | 717975 | 31810 | 5864 |
|  | Mean | 1335 | 5.94 | 680585 | 661729 | 19975 | 3794 |

### Comparison of read length retention

To test SNAILS’ efficiency, long read retention of pipetting by manual aspiration, manual rotation, and automated rotation performed by SNAILS was compared through various metrics. We observed that SNAILS produced more total reads than either manual pipetting method (Figure 2A). Across six flow cells, SNAILS produced 5,251,774 reads in total, exceeding both manual aspiration that produced 4,216,634 total reads and manual rotation that produced 2,810,324 total reads (Supplementary Table 1). The total reads generated by each flow cell is subject to many confounding variables. For instance, the number of available pores per flowcell will clearly influence sequence yield. To account for flow cell quality, we analyzed the mean reads per pore for each pipetting method (Figure 2B). SNAILS produced a mean reads per pore of 624, exceeding the mean of 543 reads sequenced by manual aspiration and the mean of 459 reads sequenced by manual rotation. However, this increase in mean reads per pore is mostly the result of DNA isolation two, where SNAILS produced a far greater number of reads than from other isolations.

**Figure 2.**
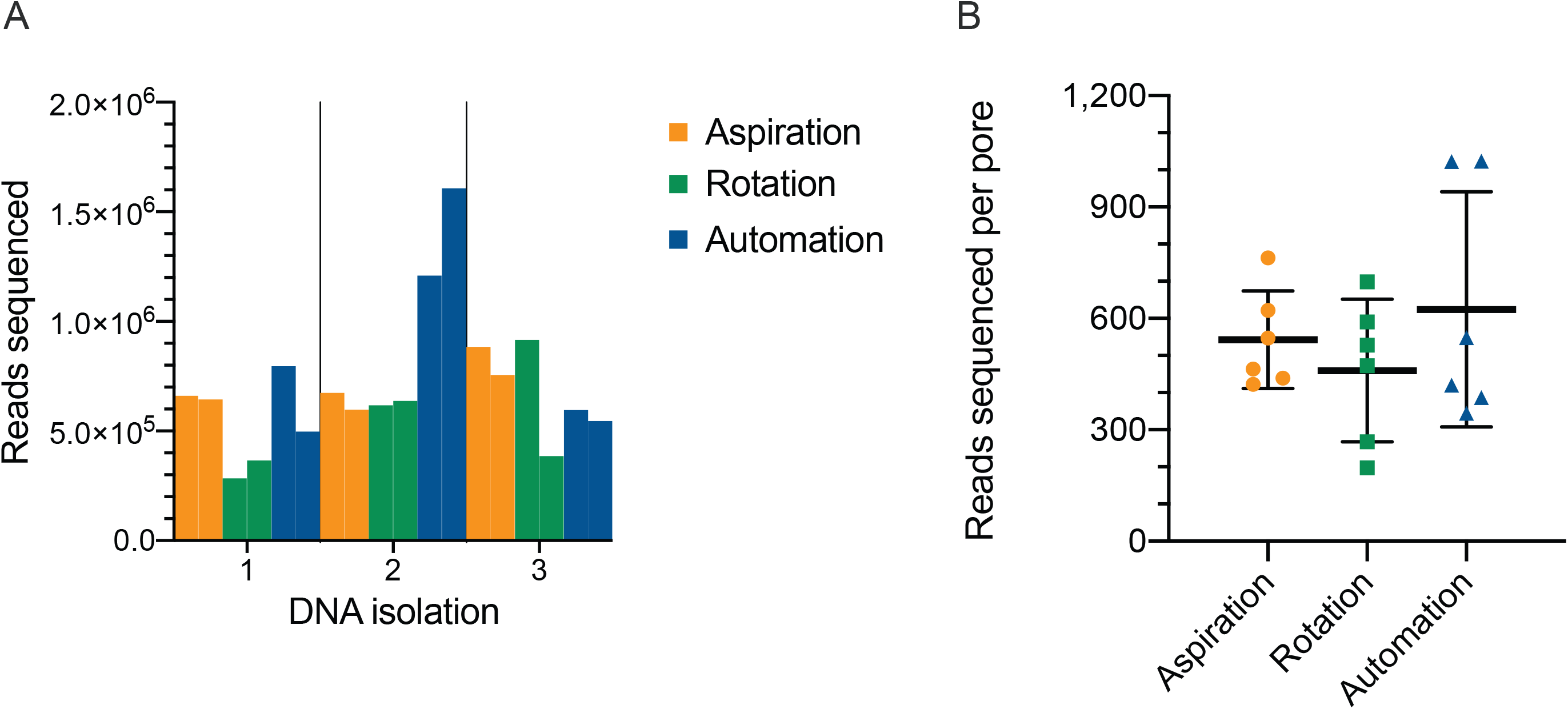
Total reads sequenced per flow cell. **A.** Total reads sequenced per library grouped by DNA isolation. **B.** Comparison of reads sequenced per available pore by pipetting method. Reads per pore we estimated by dividing the total reads per flow cell by the number of available pores at the beginning of sequencing. Available pores were calculated by MinKnow. The mean reads per pore of each pipetting method are denoted by the middle bars. The upper and lower bounds denote standard deviation.

Across eighteen library preparations, only SNAILS retained ultra-long reads in excess of one Mb: 1,253,204 bp and 1,021,308 bp (Table 2). In contrast, the maximum lengths of reads preserved by manual aspiration and rotation were 798,240 bp and 862,322 bp, respectively. SNAILS produced six reads longer than the maximum read produced by manual rotation and nine reads longer than the longest read produced by manual aspiration. These findings suggest that automated rotation of the pipette plunger via SNAILS is a superior method of read length retention though more comparative library preparations are necessary to confirm the observed trend.

**Table 2.** Ten longest mapped reads sequenced per pipetting method. Length is displayed in base pairs and was calculated after programmatic fusion of split reads (see methods). Top ten ranking is taken from the combined six sequencing libraries prepared by each pipetting method.

| Rank | Aspiration | Rotation | Automation |
| --- | --- | --- | --- |
| 1 | 798240 | 862322 | 1253204 |
| 2 | 774556 | 831285 | 1021308 |
| 3 | 737462 | 782247 | 981305 |
| 4 | 716306 | 775581 | 929976 |
| 5 | 715053 | 735123 | 907341 |
| 6 | 696967 | 729766 | 903921 |
| 7 | 694147 | 719004 | 862183 |
| 8 | 682744 | 712077 | 845167 |
| 9 | 645672 | 697032 | 800208 |
| 10 | 637514 | 673953 | 769538 |

The length distribution for the combined reads from each pipetting method is displayed in Figure 3A. Manual rotation produced a mean read length of 8,583 bp across six libraries. This surpassed SNAILS’ mean length of 6,630 bp and manual aspiration’s mean length of 5,209 bp. This hierarchy was mirrored when comparing N50s produced by each method (Figure 3C). Pipetting by manual aspiration resulted in the lowest mean N50 of 16,203 bp across six cells. By comparison, the mean N50 produced by manual and automated rotation of the plunger were 22,221 bp and 20,381 bp, respectively. SNAILS produced higher N50s in five of the six sequencing runs from the same DNA isolations over manual aspiration (Table 1). Additionally, SNAILS produced the highest N50 for a single run (31,810 bp) despite having a lower mean N50 than manual rotation (Table 1).

**Figure 3.**
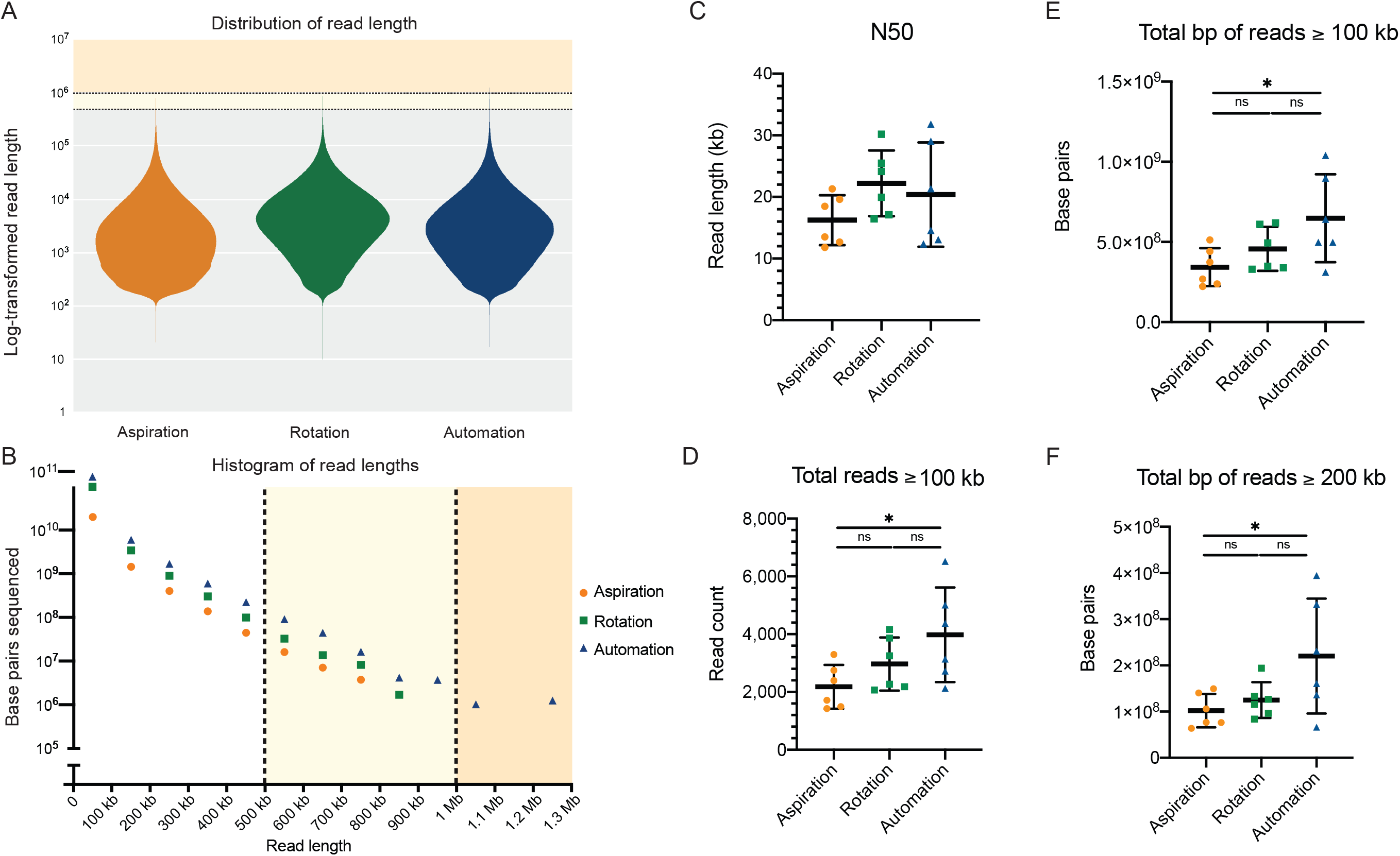
Comparison long read retention by pipetting methods. **A**. Distribution of log-transformed read lengths. Each violin plot represents the total reads sequenced across six library preparations. The Yellow region denotes 500-1000 kb. The orange region denotes 1-10 Mb **B.** Comparative read length histogram. Each point represents total base pairs sequenced across six libraries. The Y-axis denotes log-transformed total base pairs sequenced. The X-axis denotes length in base pairs. **C.** Averages of N50 values per pipetting method. Mean is denoted by center line. Standard deviation is denoted by upper and lower bounds. The Yellow region denotes 500-1000 kb, and the orange region reads greater than one Mb. **D.** Number of reads greater than 100 kb sequenced per flow cell. Significant difference is denoted by the asterisk (p-value = 0.0438). **E.** Total base pairs of reads greater than 100 kb sequenced per flow cell. Significant difference is denoted by the asterisk (p-value = 0.0415). **F.** Total base pairs of reads greater than 200 kb sequenced per flow cell. Significant difference is denoted by the asterisk (p-value = 0.0441).

When the results of all six flow cells were combined, SNAILS retained more reads of longer length than either manual method (Figure 3B). In silico modeling has predicted that an N50 greater than 100 kb will substantially improve the continuity of genomic assemblies [24]. We therefore compared the number of reads greater than or equal to 100 kb retained by each pipetting method (Figure 3D). SNAILS produced a combined 23,884 reads greater than or equal to 100 kb in length (Supplementary Table 1). This number exceeded both manual aspiration and rotation that produced 13,079 and 14,424 reads, respectively. We then compared the mean number of reads greater than or equal to 100 kb sequenced across libraries to assess whether SNAILS was more consistently retaining beneficially long reads. SNAILS produced the highest mean of 3,981. Manual rotation resulted in a mean of 2,968 and aspiration produced a mean of 2,180. The difference in means was significant between automated and aspirated pipetting by one-way ANOVA (p-value = 0.0428). No significant difference was observed between automated and manual rotation. This significant difference between manual aspiration and SNAILS was also observed by comparing the mean total base pairs of reads greater than 100 kb and 200 kb. These comparisons resulted in p-values of 0.0415 and 0.0441, respectively (Figure 3E,F). These findings suggest that SNAILS is comparable to manual rotation of the pipette plunger for length preservation through RAD004 library preparations. Furthermore, SNAILS appears to be preserving significantly more long reads than the common practice of slowly aspirating the pipette.

The initial eighteen flowcells used to compare pipetting methods resulted in relatively low N50s compared to what has been reported in the field [24]. We suspected that this may be the result of low-quality DNA isolations because all pipetting methods resulted in similarly small N50s. We developed SNAILS to increase the throughput of our internal whole genome sequencing efforts. To date, we have used SNAILS to prepare over 200 long-read libraries, some of which have resulted in N50s that exceed 100 kb. We have included sequencing results from four exemplary flow cells to demonstrate SNAILS’ ability to preserve desirably high N50s (Figure 4). These four libraries were prepared individually from four separate HMW phenol-chloroform DNA isolations. Libraries four and five were prepared from a single rhesus macaque and libraires six and seven were prepared from a single cynomolgus macaque. Each library yielded more than twenty-six thousand reads (Figure 4A). However, the first two libraries produced notably more reads when compared to the second two. This difference may be the result of less available pores at the beginning of sequencing. Flow cells four and five began sequencing with a reported 1382, and 1383 available pores whereas flow cells six and seven began sequencing with 1117 and 962 pores. The difference between runs was less pronounced when total reads were adjusted to account for available pores though still noticeably different (Figure 4B). The four flow cells resulted in N50s of 119.3 kb, 118.2 kb, 178.4 kb, and 164.5 kb, respectively (Figure 4C). Libraries five and six produced exceptionally high N50s exceeding 150 kb. It is possible that the decreased yield from flow cells five and six was due in part to the larger relative length within the libraries. We have begun investigating whether decreased yield can be partially alleviated by performing routine nuclease flushes and reloading fresh library. The ten longest mapping reads from libraries four and five all exceeded 1 Mb with a maximum length of 1.87 Mb (Supplementary Table 2). By comparison, libraries six and seven did not contain any reads exceeding 1Mb in length despite resulting in considerable larger N50s than libraries four and five.

**Figure 4.**
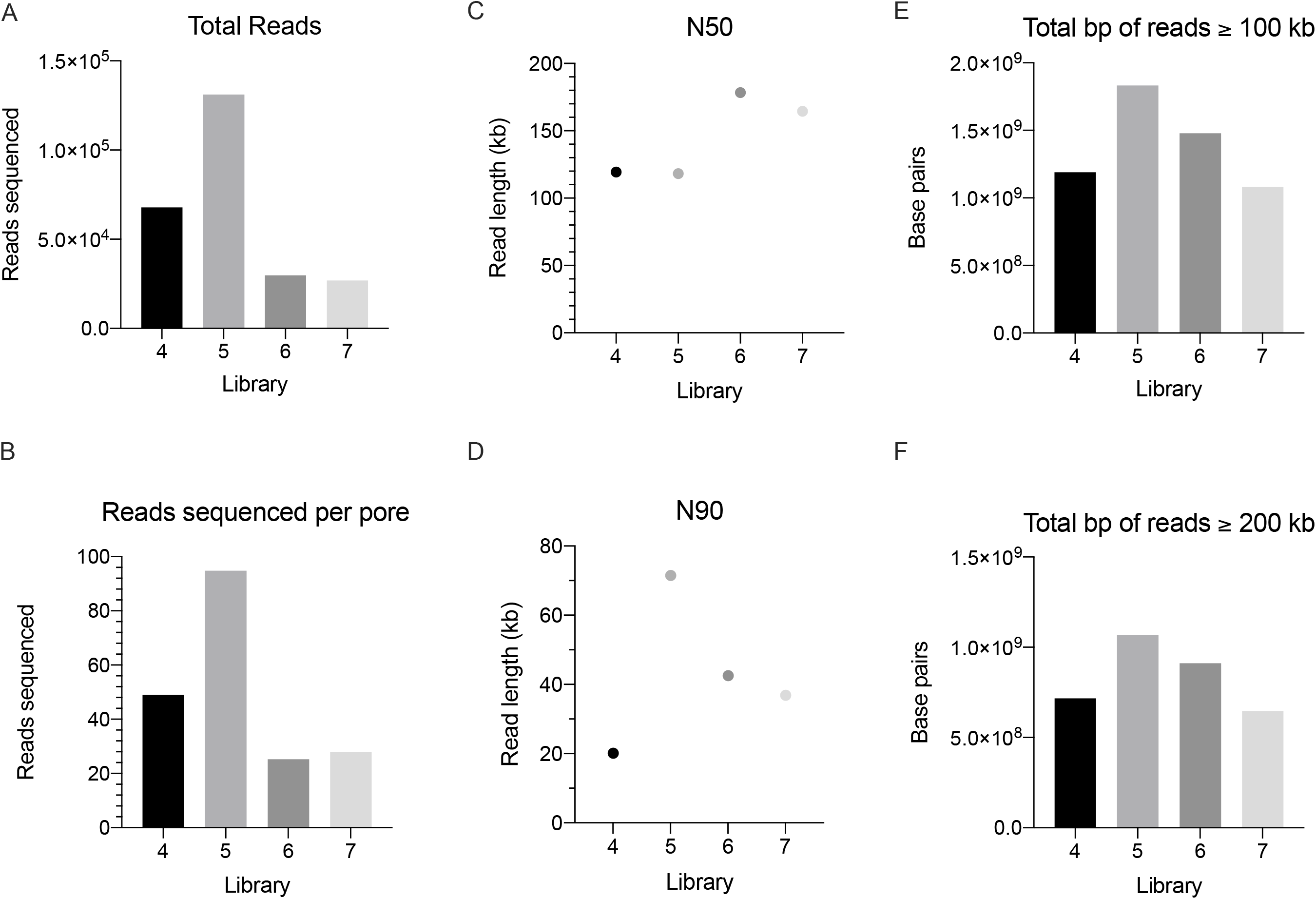
Sequencing results for four exemplary RAD004 libraries prepared by SNAILS. **A.** Total sequencing reads generated per flow cell. **B.** Reads sequenced per available pore. Reads per pore we estimated by dividing the total reads per flow cell by the number of available pores at the beginning of sequencing. Available pores were calculated by MinKnow. **C.** N50 values for each flow cell. **D.** N90 values for each flow cell. **E.** Total base pairs of reads greater than 100 kb sequenced per flow cell. **F.** Total base pairs of reads greater than 200 kb sequenced per flow cell.

All four runs resulted in over a combined 1.08 Gigabase pairs from reads greater than 100 kb in length and a combined 647 Mb from reads greater than 200 kb in length (Figure 4 E,F). By comparison, the longest library prepared by SNAILS from the initial eighteen libraries resulted in a combined 1.04 Gigabase pairs from reads greater than 100 kb in length and 332 Mb from reads greater than 200 kb in length (Figure 3E, F). This further suggests that the quality of HMW DNA isolation is perhaps a larger determinant of relative DNA length given that SNAILS were used to prepare all five libraries. Based on this theory, we are actively working to adapt DNA purification kits manufactured by Circulomics and New England Biolabs into our workflows. Our thought is that phenol-chloroform DNA purification may be difficult to perform reliably and that perhaps an alternative purification methodology can more consistently produce desirable HMW DNA. Library five appeared to be somewhat exceptional in both yield and length. This library resulted in an N90 of 71.5 kb meaning 90% of the reads in the library were 71.5 kb or greater in length (Figure 4D). These results are somewhat perplexing given that library five was prepared identically to the other presented libraries and began sequencing with nearly the same amount available pores as library four (1382, and 1383, respectively). It therefor stands that additional factors play an important roll in the determination of DNA length through sequencing. However, the results of the presented four exemplary libraries demonstrate that SNAILS’ slow pipetting is capable of maintaining N50s of desirable length with the obvious advantage of alleviating time-consuming pipetting while increasing throughput.

## Discussion

Minimizing HMW DNA shearing during library preparation is a necessary precaution needed to maximize the advantages of ONT long read sequencing. We have designed and tested SNAILS, a 3D-printed instrument that automates the slow pipetting steps used during the modified RAD004 library preparation protocol. SNAILS retain more long reads when compared to libraries prepared by conventional slow aspiration pipetting. The amount of manual preparation time saved is arguably the most considerable advantage provided by SNAILS. The RAD004 long read protocol typically requires at least two hours to prepare a single library for only eighteen pipetting steps [13]. Developing the ability to prepare long read libraries requires substantial training to execute properly. Moreover, only technicians with steady and dexterous hands can effectively pipette at the consistent, slow rate necessary to preserve length. SNAILS reduces the amount of hands-on time required to prepare libraries to a few minutes and the technical burden to executing a couple of commands. This enables task-shifting to less trained staff. Moving forward, we plan to expand on the SNAILS design by creating a pipette-mounted version. A pipette-mounted chassis could be attached to a poseable or robotic arm to give SNAILS 360 degrees of motion. This design would essentially eliminate any manual intervention that remains when using the present SNAILS design, thereby completely automating long read library preparation via the RAD004 protocol. Furthermore, increased range of motion could allow SNAILS to be more easily applied to a broader range of protocols requiring delicate pipetting of samples such as Circulomics’ disk-based or phenol-chloroform extractions. John Tyson’s Rocky Mountain protocol is a promising approach to increasing N50s using the ligation sequencing kit and forgoing bead cleanups. The anticipated Circulomics’ ultra-long library preparation kits are also reporting encouraging early results. It is our understanding that all these methods require delicate pipetting of samples and reagents. Therefore, SNAILS can be adapted to fit the protocols’ needs and eliminate the risk of shearing caused by manual pipetting.

Liquid handling robots manufactured by Tecan, Eppendorf, and others are theoretically capable of automating long read library preparation by designing custom protocols. However, liquid handling robots are not designed for slow pipetting and require knowledge of the instrument’s programmatic background to implement custom protocols. By comparison, SNAILS is explicitly designed for slow pipetting and operates with pythonic code that requires little programmatic expertise to customize the speed, draw volume, and number of pipetting steps. This specialization sacrifices SNAILS’ ability to perform more complex protocols. However, SNAILS has a meager cost of entry when compared to enterprise liquid handling robots. 3D printing has become an increasingly utilized resource in the development of biological instrumentation and laboratory consumables [27]. In the past five years, 3D printers have rapidly decreased in price, fostering an open-sourced, at-home printing community. At the writing of this manuscript, Creatality 3D’s Ender 3 printer can be purchased for under $200 USD and is equipped with a bed large enough to handle printing pieces for SNAILS. When printed using 20% infill and support density of 25, the entire structure amounts to roughly 350 grams of polylactic acid, which costs approximately $30 USD. The remaining electronic components can be inexpensively sourced (Supplementary Table 2). Together, the total cost of a single SNAILS is roughly $165 USD. This cost of entry pales in comparison to the cost of labor required for the same task. Because of this entry cost, multiple SNAILS can be assembled and run in tandem by a single technician. For example, we have deployed SNAILS that are often run simultaneously to produce eight long read libraries in a single day. The SNAILS approach to library preparation is an economically viable solution for increasing throughput without a significant sacrifice to read length.

Manual slow pipetting is inherently difficult to perform. Maintaining a consistent volumetric flow rate isn’t easily measurable or reproducible when slow pipetting is performed by hand. Minute changes in volumetric flow rate caused by slight hand movements can result in DNA shearing. SNAILS resolves these inconsistencies through its voltage-dependent volumetric flow rate. The voltage supplied to SNAILS’ motor can be manipulated using a pulse width modulation signal delivered by a microcontroller to the L298N driver. The stepwise changes in pipetting rate can be accurately benchmarked to optimize ultra-long read retention. Any pipetting will inevitably result in DNA shearing, yet performing multiple pipetting steps is necessary to ligate sequencing adaptors to DNA molecules. Theoretically, performing less pipetting steps in library preparation will result in less DNA shearing. However, this may come at the cost of total reads sequenced. SNAILS can help establish optimal protocol conditions to maximize length and long read yield.

As presented here, SNAILS are an admittedly imperfect system with several remaining areas of improvement. Though SNAILS’ speed of pipetting seems to accommodate highly viscous HMW DNA, DNA shearing still may occur. Across six flow cells, SNAILS showed larger variation in N50 than manual rotation of the pipette. This may be a result of technical ability as manual rotation was performed by experienced technician while SNAILS were operated by a more novice technician. The total yield of each flow could possibly have been increased by implementing nuclease flushes. After the preparation of this manuscript, we have used nuclease flushes with SNAILS, which has resulted in a general increase in throughput. However, more factors are involved in maintaining long sequences than the rate and consistency of pipetting. A more in-depth investigation into the factors affecting length retention is necessary to help guide the development of SNAILS and other ultra-long read protocols.

## Conclusions

SNAILS is a cost-effective, easily-deployable, robotic pipette that automates long read RAD004 library preparation for ONT sequencing. SNAILS outperforms conventional slow pipetting in terms of preserving reads exceeding 100 kb in length and acquiring larger N50s within a sequencing library. Additionally, SNAILS allow untrained technicians to perform effective slow pipetting while simultaneously increasing throughput of a single technician without sacrificing the libraries quality. SNAILS serves as a platform for immediately increasing the quantity and quality of long read libraries and provides a basis for future optimization of long read retention.

## Methods

### CAD, 3D printing

All 3D printed portions of the robot were designed in Solidworks 2019 (https://www.solidworks.com/), a computer-aided designing and computer-aided engineering program. All 3D printing was conducted on Ultimaker 3, S3, and S5 machines with polylactic acid. Solidworks part files were converted to STL files and imported to the Ultimaker Cura program (https://github.com/Ultimaker/Cura). Parts were positioned on the build plate at least 0.5 cm apart with flat faces down. For the towers, the layer height was set to 0.2 mm, infill to 20%, support material unchecked, and the remaining settings left at default. For the base, motor holder, and bridge parts, the layer height was set at 0.2 mm, infill to 20%, support density to 25%, support material checked, and the remaining settings left at default. Before submitting the print job to the printer, the Ultimaker was loaded with polylactic acid and the build plate was covered in glue from the Ultimaker glue sticks to ensure the parts would not shift while printing. The Solidworks parts were converted to Solidworks drawings to create the figures. The *e - landscape.slddrt* size sheet was used as a base, and the sheet format was suppressed. Multiple angles for each part were included using the Model View tool under View Layout and dimensions were added using the Smart Dimensions tool under Annotation with the significant figures, size, and units edited in the Document Properties under Dimensions.

### SNAILS assembly

All support material was removed, and dovetail joints were assembled. Due to the margin of error in printing using the Ultimaker, a Dremel was used to shave down parts and a mallet was used to squarely mesh dovetail joints. Raspbian v4.19 was installed using NOOBS v3.3.1 to a 32 GB micro sd card and booted on a Raspberry Pi model 3B+. Electronic components were assembled into a solderless circuit as depicted in Figure 1C. Briefly, the general-purpose input/output pins 23, 24, 25, and ground pin 6 were connected to IN1, IN2, ENA, and GND terminals of the L298N motor driver board, respectively. The positive and negative leads of a twelve-volt direct current power supply were connected to the twelve-volt and GND terminals of the L298N motor driver, respectively. Finally, the two terminals of a twelve-volt direct current planetary motor were connected to the OUT1 and OUT2 terminals of the L298N motor driver. Protocols were written in python3 and downloaded to the rpi (https://github.com/emmakn/SNAILS). All sources for parts used in assembly are displayed in Supplementary Table 3..

### Animal selection and sample collection

Blood was obtained from two rhesus macaques (rh1990, r02072) and one cynomolgus macaque (cy0161) housed at the Wisconsin National Primate Research Center. The animals were released after blood was collected. Peripheral blood mononuclear cells were isolated from the buffy coat following density gradient centrifugation with ficoll and frozen in liquid nitrogen. Sampling was performed following protocols approved by the University of Wisconsin-Madison Institutional Animal Care and Use Committee and in accordance with the regulations and guidelines outlined in the Animal Welfare Act, the Guide for the Care and Use of Laboratory Animals, and the Weatherall report (https://mrc.ukri.org/documents/pdf/the-use-of-non-human-primates-in-research/).

### High Molecular Weight DNA Extraction

DNA was extracted from frozen peripheral blood mononuclear cells using a previously described ultra-long DNA extraction protocol [24] with some modifications. For each extraction, roughly twenty million cells were thawed and then spun at 300 x g for 10 min. The supernatant was removed, and the pelleted cells were resuspended in 200 μl PBS. In a 15 ml conical tube, the resuspended cells were added to 10 ml of TLB (100 mM NaCl, 10 mM Tris-Cl pH 8.0, 25 mM EDTA pH 8.0, 0.5% (w/v) SDS, 20 ug/ml Qiagen RNase A, H_2_O) and vortexed at full speed for 5 sec. The samples were then incubated at 37°C for 1 hr. Qiagen Proteinase K (200 ug/ml) was then added and mixed by slow end-over-end inversion three times. Samples were incubated at 50°C for 2 hrs with 3x end-over-end inversion every 30 min. The lysate was slowly pipetted into two phase-lock 15 ml conical tubes in 5 ml increments. The phase lock conical tubes were prepared prior to DNA isolation by adding ~2 ml autoclaved high-vacuum silicone grease into 15 ml conical tubes and spinning max speed for 1 min. To each phase-lock tube of lysate, 2.5 ml buffer-saturated phenol and 2.5 ml chloroform were added before rotational mixing at 20 rpm for 10 min. Tubes were then spun down at 4000 rpm for 10 min in an Eppendorf 5810R centrifuge with a swinging-bucket rotor. After centrifugation, the aqueous phase was poured slowly into a new phase-lock 15 ml conical tube. The addition of phenol and chloroform, rotational mixing, and centrifugation was repeated with the second set of phase-lock tubes. The aqueous phases from both phase-lock tubes were slowly poured into a single 50 ml conical tube before adding 4 ml of 5 M ammonium acetate and 30 ml of ice-cold 100% ethanol. The mixture was incubated at room temperature while the DNA visibly precipitated. Once the DNA rose to the surface of the mixture, a glass capillary hook was used to retrieve the DNA. The hooked DNA was dipped into 70% ethanol, then carefully worked off of the hook into an Eppendorf DNA LoBind 1.5 ml tube. One ml of 70% ethanol was added to the tube before spinning down at 10,000 x g for 1 min and the supernatant was removed. This process was repeated for a second 70% ethanol wash, and any remaining ethanol was evaporated off during a 15 min incubation at room temperature. 100 μl of elution buffer (10 mM Tris-Cl pH 8.0, H_2_O) was added before incubating the DNA at 4°C for at least two days. A total of three ultra-long DNA isolations were performed.

### Library Preparation

For each library preparation, pipetting was performed using one of three techniques: manual aspiration, manual rotation, and automated rotation with SNAILS. All pipetting was performed with wide-bore pipette tips. For manual aspiration libraries, volumes were pipetted up and down by hand as slowly as possible. For manual rotation, the pipette plunger was twisted by hand as slowly as possible to aspirate and dispense. For automated rotation, a SNAILS was programmed to rotate the pipette plunger at a rate of approximately 1.25 rotations per minute. An aliquot of 16 μl HMW DNA from each library was loaded into a flow cell on an Oxford Nanopore Technologies (Oxford, United Kingdom) GridION. We used the ONT SQK-RAD004 Rapid Sequencing Kit with modifications of a previously described library preparation [24] optimized for this kit. This procedure is also described in Karl et al. (manuscript in publication). 1.5 μl fragmentation mix (FRA) and 3.5 μl of elution buffer were added to the DNA aliquot and pipetted five times to mix. The samples were then placed on an Applied Biosystems Thermal Cycler (ThermoFisher Scientific, Waltham, MA, USA) at 30°C for 1 min followed by 80°C for 1 min. One μl of rapid adaptor (RAP) was added to the solution and pipetted five times. The library was then incubated at room temperature while the flow cells were primed as follows. For each flow cell, 30 μl of flush tether (FLT) was added to a tube of flush buffer (FLB), and a very small volume of buffer was removed from the priming port in order to remove the air gap. 800 μl of the Flush tether + Flush buffer solution were added to the priming port. Following a five minute incubation, the SpotOn port was opened and 200 μl of the FLT + FLB solution was slowly added to the priming port so that small volumes of solution rise from the SpotOn port and then return to the cell. 34 μl of the sequencing buffer (SQB) and 20 μl of water were added to the sample solution and pipetted three times. Finally, 75 μl of the prepared library was slowly drawn into a pipette with a wide bore tip. The library was then added dropwise to the SpotON port.

### MinION Sequencing and base calling

Eighteen R9.4 (FLO-MIN106) total MinION flow cells were loaded with ultra-long DNA libraries and sequenced for 48 hours and default parameters on a GridION instrument according to ONT guidelines using their MinKNOW software to control each sequencing run. Different versions of MinKNOW were used throughout the course of the study as updates were released; specific versions are recorded in the metadata in the fast5 files for each run. Only reads that passed the preliminary default screening of MinKNOW were used for base calling. Base calling was performed using the GPU-enabled guppy version 3.2.4 software provided by ONT with the following settings: [--config dna_r9.4.1_450bps_hac.cfg --num_callers 14 --chunk_size 500 --gpu_runners_per_device 8 --chunks_per_runner 768]. Currently, Guppy and MinKNOW are only available to ONT customers via their community site (https://community.nanoporetech.com).

### Fusing split reads and read statistics

Whale_watch.py was used to identify and fuse reads that were incorrectly split during sequencing as previously described [26]. Briefly, basecalled fastq files were merged into a single file and mapped against the human reference genome GRCh38 [28] using minimap2 [29] and saved as a .paf output file. Next, whale_merge.py was run on the resulting .paf files along with their corresponding merged fastq files to output fastq files containing fused reads. Read lengths, N50s, and N90s for fused fastq files were calculated using stats.py from the bbtools suite (https://sourceforge.net/projects/bbmap/). Figures and statistics were generated using Prism 8.4.0 for MacOS (GraphPad Software, San Diego, California USA, www.graphpad.com).

## Abbreviations

SNAILS: Slow nucleic acid instrument for long sequences
RAD: Rapid adaptor
ONT: Oxford Nanopore Technologies
HMW: high molecular weight
bp: base pairs
kb: kilobase pair
Mb: Megabase pair

## Declarations

### Ethical approval and consent to participate

The rhesus macaque used in this study was cared for by the staff at the Wisconsin National Primate Research Center in accordance with the regulations, guidelines, and recommendations outlined in the Animal Welfare Act, the Guide for the Care and Use of Laboratory Animals, and the Weatherall report (55–57). The University of Wisconsin-Madison College of Letters and Science and Vice Chancellor for Research and Graduate Education Centers Institutional Animal Care and Use Committee approved the nonhuman primate research covered under protocol G005401-R01. The University of Wisconsin-Madison Institutional Biosafety Committee approved this work under protocol B00000117.

### Consent for publication

Not applicable.

### Availability of data and materials

The datasets used during the current study are available from the corresponding author on reasonable request. The accompanying files and schematics are freely available for use at https://github.com/dholab/SNAILS.

### Authors’ contributions

TP designed and supervised the project, performed the analysis, and wrote the manuscript. TP and EN created the SNAILS design. EN created CAD schematics, 3D printed designs and wrote the manuscript. TP, JK, EN, and CS performed sequencing experiments. DB wrote software code. HB, RW, and DO provided data interpretation. All authors read and approved the final manuscript.

## Acknowledgements

The authors would like to thank Phoenix Shepherd for creating the SNAILS acronym.

