## Supplementary figures and images for "Consistent ultra-long DNA sequencing with automated slow pipetting"

### Supplemental Figure 1

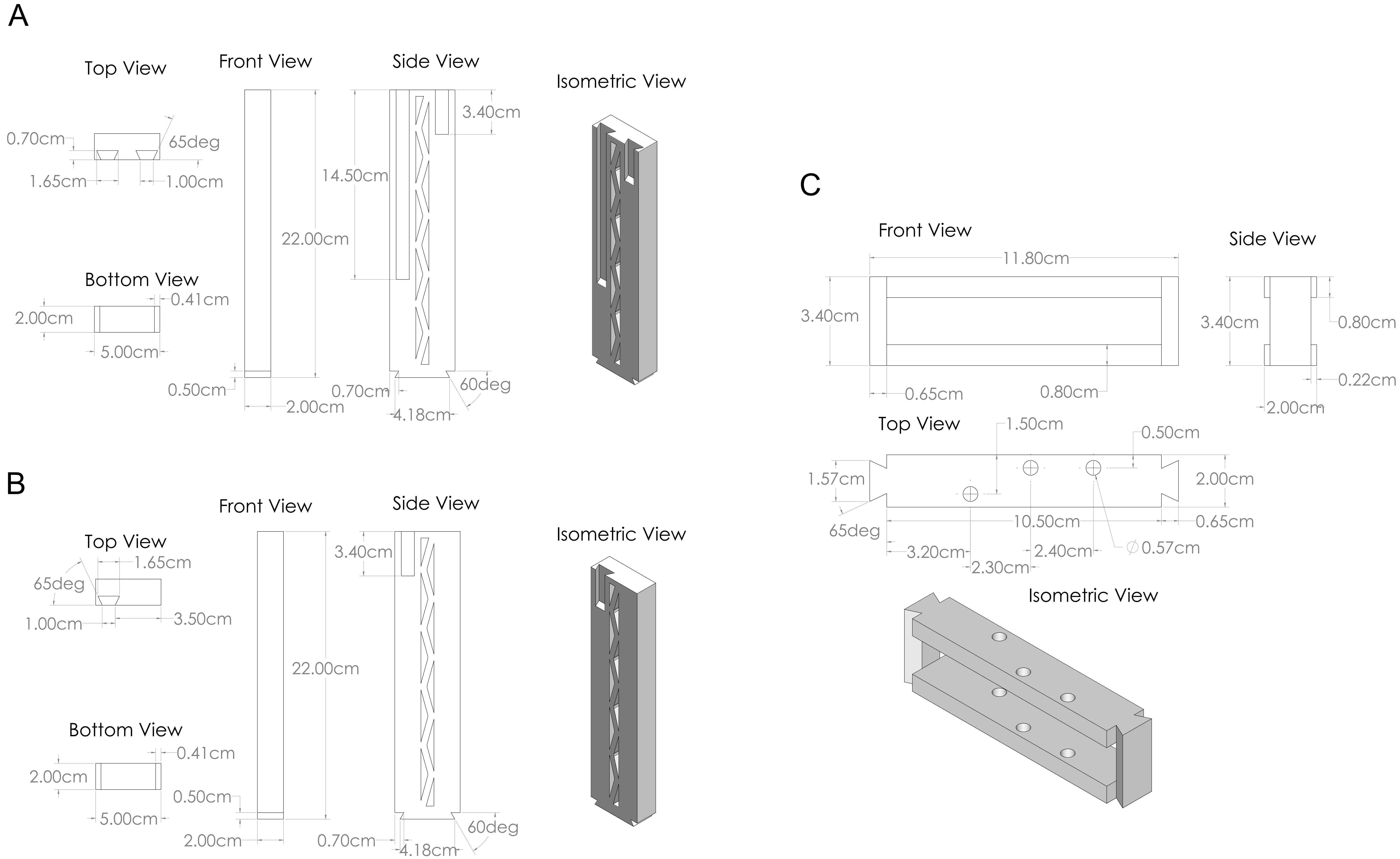

### Supplemental Figure 2

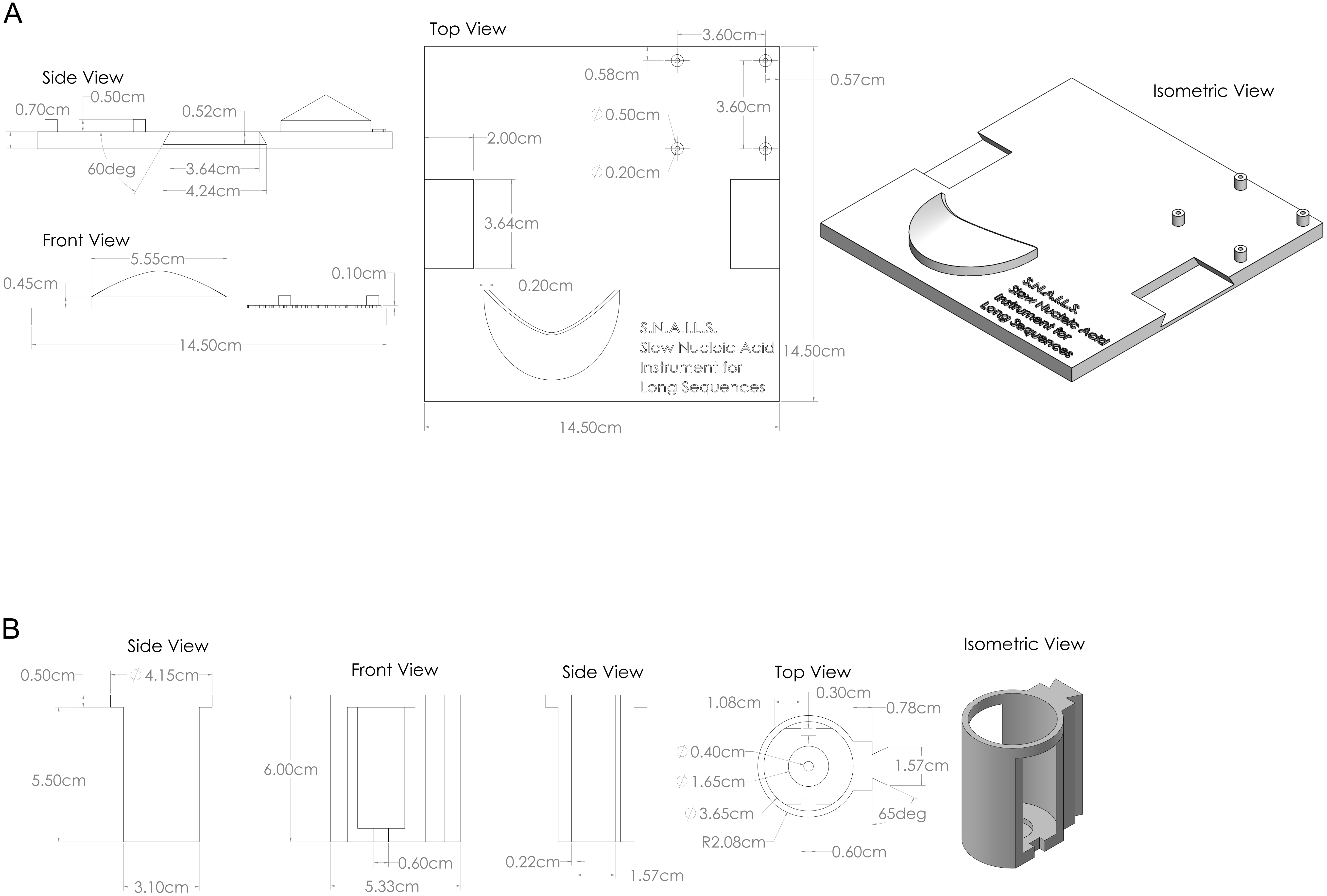

### Supplemental Figure 3

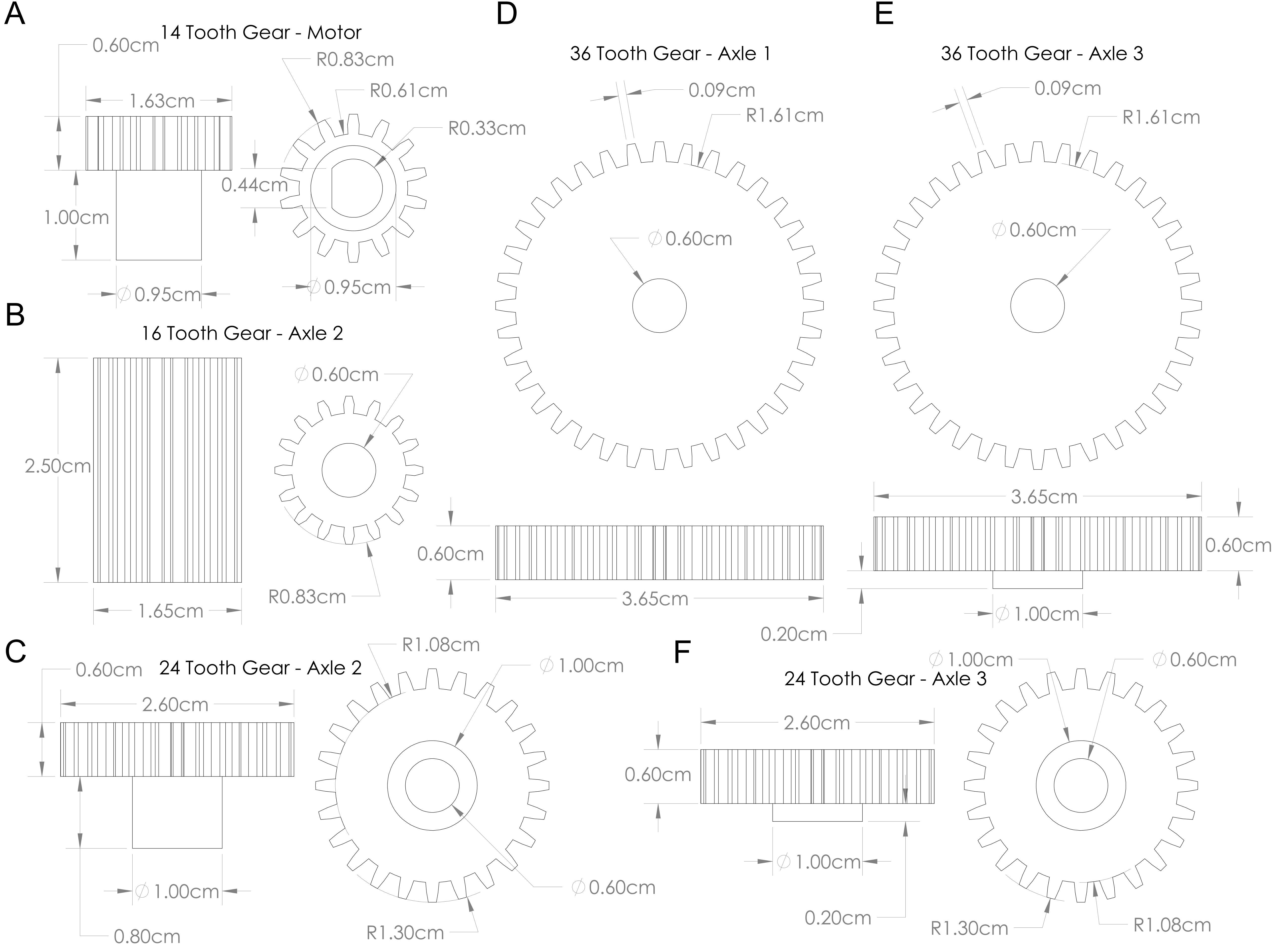

### Supplemental Figure 4

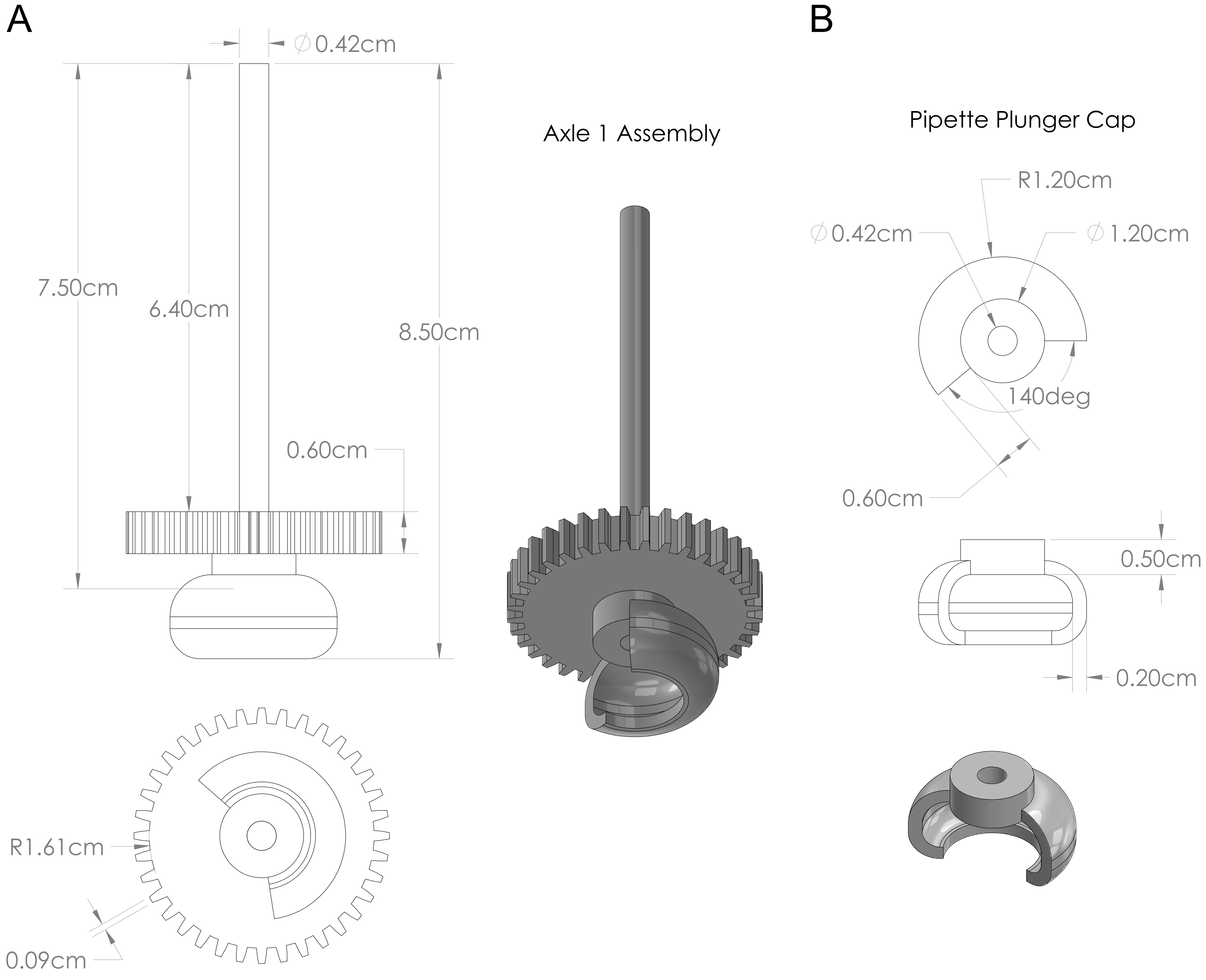
