## Supplemental Information for "Consistent ultra-long DNA sequencing with automated slow pipetting"

**Supplementary Figure 1. Schematics of SNAILS towers and bridge.** **A.** Dimensions of the right, dovetailed tower. **B.** Dimensions of the left, dovetailed tower. **C.** Dimensions of the dovetailed bridge which houses gear axles.

**Supplementary Figure 2. Schematics of SNAILS base and motor basket. A.** Dimensions of the base. The slots on either side of the base accept each towers’ dovetail joints. The base is designed with a protruding slot that fits around the base of an Ovation M micropipette. The back right corner includes standoffs aligned to L298N boards for attachment. **B.** Dimensions of the dovetailed motor basket. The basket is fitted to a 12 rpm HD premium planetary gear motor.

**Supplementary Figure 3. Schematics of SNAILS gears.** **A.** Dimensions of 14 tooth gear cap for planetary motor. **B.** Dimensions for 16 tooth axle 2 gear. **C.** Dimensions of 24 tooth axle 2 gear. **D.** Dimensions of 36 tooth axle 1 gear. **E.** Dimension of 36 tooth axle 3 gear. **F.** Dimensions of 24 tooth axle 3 gear.

**Supplementary Figure 4. Schematics of SNAILS axle 1 and pipette cap**. **A.** Dimensions for axle 1. **B.** Dimensions for pipette cap. The cap is fitted for an Ovation M Micropipette.

**Supplementary Figure 5: Sequencing results for three exemplary RAD004 libraries prepared by SNAILS.** **A.** Total reads sequenced per flow cell. **B.** N50 per flow cell. **C.** Total basepairs of reads greater than or equal to 100 kb per flow cell. **D.** Total base pairs of reads greater than or equal to 200 kb per flow cell

**Supplementary Table 1. Combined number of reads sequenced by length.** Numbers are indicative of the combined reads sequenced from six separate library preparations.

**Supplementary Table 2. Key resources.** Vendors and product IDs are listen where applicable.

**Supplementary Table 3. Top ten longest mapped reads sequenced per pipetting method.** Length is displayed in base pairs and was calculated after programmatic fusion of split reads (see methods).
